# LAMPrEY: a Python-based automated quality control tool for large-scale proteomics datasets

**DOI:** 10.64898/2026.05.06.722826

**Authors:** Mario E. Valdés-Tresanco, Soren Wacker, Mario S. Valdés-Tresanco, Andriy Plakhotnyk, Nicholas I. Brodie, Morgan Hepburn, Annegret Ulke-Lemée, Edward L. Huttlin, Ian A. Lewis

## Abstract

Over the past years, proteomics has moved increasingly towards the analysis of large cohorts of biological specimens. This has been made possible by significant improvements in mass spectrometry technology, chromatographic separation methods, and improved data acquisition strategies. These technological advances now routinely enable experiments that yield vast datasets that substantially outstrip the capacity of existing proteomics data analysis approaches. Processing such large datasets requires purpose-built, quality control tools designed to organize and analyze the data while recording all processing parameters for reproducibility. To address this need, we developed an open-source, Python-based software platform, **L**arge-scale **A**utomated **M**ulti-level **Pr**oteomics **E**valuation by P**y**thon (LAMPrEY), a comprehensive quality-control pipeline for quantitative proteomics analyses of large cohorts of samples. LAMPrEY features GUI-based file submission, automated processing with MaxQuant and RawTools, an interactive analytics dashboard, and an application programming interface (API) for programmatic usage that collectively enable rapid, reproducible analysis and interpretation of proteomics data. We demonstrate the longitudinal monitoring and analytical capabilities of LAMPrEY using TMT11 quantitative proteomics data generated from 910 *Enterococcus faecium* isolates collected from bloodstream infection patients. LAMPrEY is an open-source software that can be accessed at www.lewisresearchgroup.org/software.

## INTRODUCTION

Proteomics based on liquid-chromatography mass-spectrometry (LC-MS) has emerged as a mainstream analytical tool for studying complex biological systems and understanding the response of organisms to genetic, environmental, or nutritional perturbations.^1–3^ Over the last decade, researchers have increasingly applied proteomics to large-cohort studies with hundreds to thousands of samples.^4–9^ Major progress has been enabled through advances in hardware, such as the chromatography platform EvoSep,^10,11^ mass-spectrometry platforms such as the Thermo-Fisher Lumos,^12^ Orbitrap Astral,^13^ and the Bruker timsTOF Ultra 2.^14,15^ Moreover, methodological refinements such as increasingly powerful data-independent acquisition strategies and higher-multiplex chemical labeling systems such as TMTpro 35-plex have facilitated the analysis of large cohorts of specimens.^16^ Collectively, these advances have vastly expanded the practical scope of proteomics.

Although the capacity to generate data has increased significantly over the past decade, software tools that allow users to process and track data quality across large cohorts have not kept pace with these innovations. This presents a central challenge because longitudinal proteomics datasets are vulnerable to a wide range of technical confounders, including chromatographic column aging, gradual changes in instrument performance, lot-to-lot variation in consumables, and errors introduced during sample preparation. In large experiments that require data acquisition over long timeframes, such effects accumulate and can manifest as systematic drift, ultimately reducing data quality, comparability, and confidence in downstream biological interpretation.^17,18^ Tracking data integrity over large-cohort experiments is crucial. Although there is a wide collection of programs available for proteomics data processing and quality control (QC), most tools are not designed for large-cohort experiments.^19–26^ Moreover, these tools are typically not true end-to-end applications but rather represent a single link in a chain of tools and processing steps. Such tool chains rely heavily on manual intervention, requiring significant time and effort from the user and increasing the risk of human error. To address these challenges, we developed **L**arge-scale **A**utomated **M**ulti-level **Pr**oteomics **E**valuation by P**y**thon (LAMPrEY), a Python-based application and end-to-end tool for automated, reproducible processing of proteomics data and QC analysis in large-cohort projects. LAMPrEY uses established tools (MaxQuant^27^ and RawTools^28^) to process Thermo Fisher RAW Files, and wraps the results with convenient automation, data management, and visualization layers.

## EXPERIMENTAL SECTION

### Sample Processing

Bacterial isolates used in this study were retrieved from the *Calgary BSI Cohort*, a collection of 38,000 microbes obtained from bloodstream infection patients in the Calgary, AB metropolitan health zone. The data used here were generated from 910 *Enterococcus faecium* isolates collected between 2006 and 2022. Frozen clinical *E. faecium* isolates were retrieved from Alberta Precision Laboratories (APL; Alberta’s diagnostic services provider) and were grown at 37 °C with 5% CO_2_ in a deep-well plate to logarithmic phase (optical density A_600_ ∼ 0.4) and re-frozen at -80 °C. Cultures were thawed and collected by quick centrifugation. Concentrated lysis buffer (10X) was added to cultures to achieve final concentrations of 1% w/v SDS (CAS 11-21-3), 0.5% v/v NP-40 (CAS 9036-19-5), 0.5% v/v Triton X-100 (CAS 9002-93-1), and 0.1 µg/ml BSA. Next, the cells were disrupted using ceramic beads on an OMNI Beadruptor 96 (Kennesaw, GA, US). Proteins were then isolated and cleaned using the SP3 method.^29^ Isolated proteins were digested overnight with 2 µg mass spectrometry-grade trypsin/LysC (Thermo Fisher, A40009). Finally, the peptide samples were labelled with Tandem Mass Tag (TMT11-131C Label Reagent; Thermo Fisher, A34808)^30^, quenched, and pooled along the rows to generate eight multiplexes (A to H) alongside a bridge sample. The bridge sample was prepared from a group of *E. faecium* isolates representative of the cohort’s genetic diversity and was intended to provide a common quantitative benchmark. Pooled samples were C18 cleaned and then evaporated to dryness. A Microlab® Nimbus (Hamilton Company, Reno, NV) liquid handler with 3 × 4 deck spaces and an 8-channel pipet head, 96-well plate heater shaker (3 mm shaking amplitude), CO-RE Gripper Paddles, and HEPA filtered hood equipped with a 96-well magnet (Magnum FLX, Alpaqua, Beverly, MA, US) was used for SP3, peptide normalization, TMT-labeling, and pooling of samples.

### Liquid Chromatography-Mass Spectrometry

Peptides were resuspended in nano-LC running buffer (0.1%, v/v, formic acid) and analyzed by LC-MS/MS using an Orbitrap Fusion Lumos (Thermo Fisher) coupled to an EASY-nLC1200 liquid chromatography system with an EasySpray source. Peptides were injected and separated over 90 minutes on a 15 cm column (ES801A, 2 µm bead size, 50 µm I.D.), and protected by a Pepmap 100 precolumn (164946-2, 75 µm I.D., 2cm length). Running buffers were 0.1% (v/v) formic acid and 80% (v/v) acetonitrile with 0.1% (v/v) formic acid. Eluted peptides were analyzed with a single FAIMS energy (-40CV) and MS2 scans. MS1 precursor scans were performed in the Orbitrap (mass range 375 - 1600 m/z), with an Automatic Gain Control (AGC) target of 4 × 10^5^ or 50 ms maximum injection time. Peptides with a charge stage of 2 – 6 with a minimum intensity of 25,000 were isolated in the quadrupole (isolation window 0.7) and fragmented (fixed collision energy 38%). Fragments were analyzed in the Orbitrap (resolution 5 × 10^5^, scan range 100 - 2000 m/z) with an AGC target of 25,000. Precursors were excluded for 45 seconds after a single detection.

### Data Processing and Analysis

Mass spectrometry files were processed individually using MaxQuant (v2.4.12.0) and RawTools (v2.0.3). RawTools was run with parameters “-p -q -x -u -l -m -r TMT11 -chro 12TB”. In MaxQuant, digestion was set to Trypsin and LysC. TMT tags on lysine residues and peptide N-termini were set as fixed modifications, and methionine oxidation as a variable modification. A maximum of two missed cleavage sites and peptide length of 8-25 amino acids was enforced. Quantification was achieved by summing TMT-derived peptide intensities per protein. Protein intensities were normalized as described elsewhere.^18^

## RESULTS AND DISCUSSION

### LAMPrEY Implementation

LAMPrEY was implemented in Python 3.10 using Django (v3.2). The platform can be run locally on a single workstation or deployed as a server-based application for shared laboratory access. The backend automates processing with MaxQuant and RawTools, stores the results for all the runs, and also exposes an authenticated API for programmatic use. We provide LAMPrEY as a Docker container for portability across different operating systems.

Users interact with LAMPrEY through a browser-based graphical user interface (GUI) that consists of three integrated components: 1) the *Admin* space for project and pipeline configuration, 2) the *Main* section for file submission and run tracking, and 3) the interactive *Dashboard* for data exploration (Figure 1). The *Admin* interface is password-protected and is intended for a limited number of authorized administrators (Figure S1). In this section, independent projects can be created and users assigned to each project. Each project can contain multiple analysis pipelines (Figure S2). Each pipeline stores the processing configuration for uploaded RAW files, including a protein database in FASTA format, RawTools parameters, and an optional MaxQuant parameter file (mqpar.xml). Pipelines can be configured to use different MaxQuant versions and different mqpar.xml templates suitable for TMT-based quantitative workflows. By default, LAMPrEY includes a bundled installation of MaxQuant v2.4.12.0 together with a default mqpar.xml template for TMT11 analysis (Figure S3). The centralized control of pipeline configurations, software versions, and analysis parameters ensures that the same processing elements are applied to proteomics data generated over extended periods.

**Figure 1.**
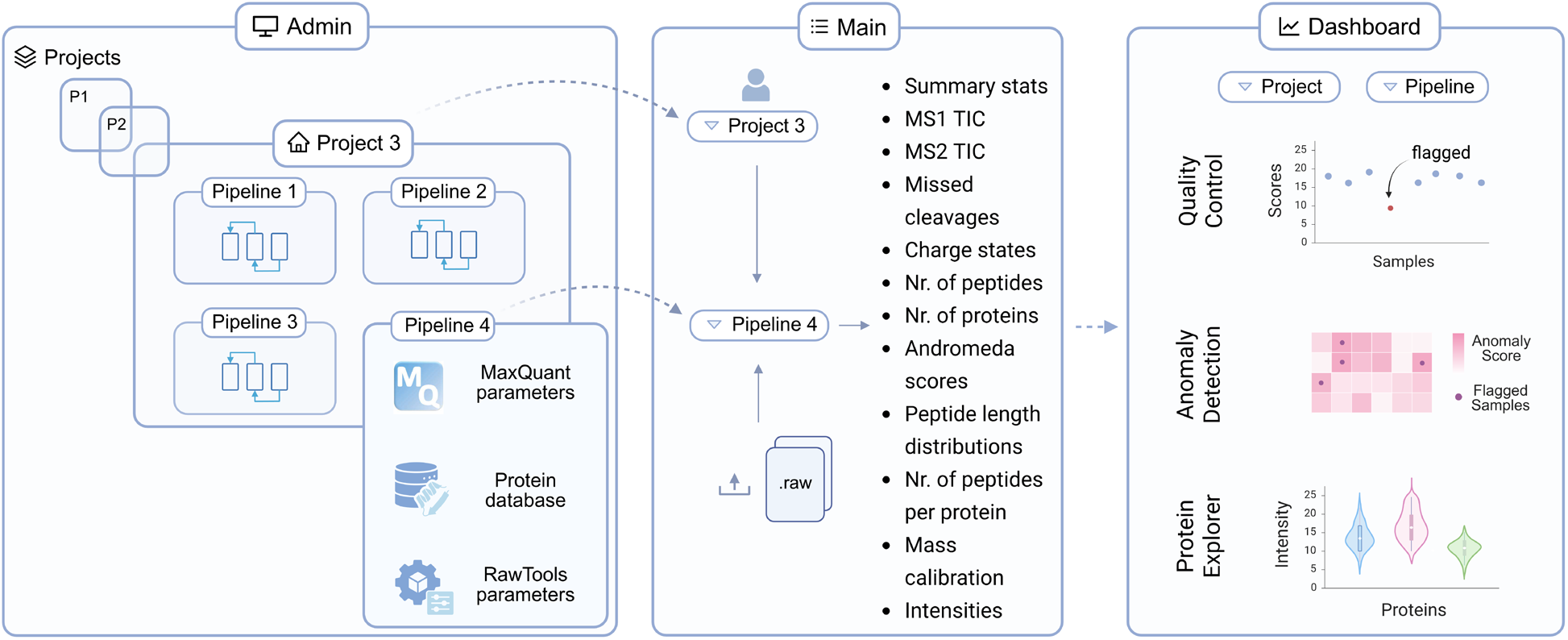
LAMPrEY components. LAMPrEY consists of three integrated components: 1) *Admin*, 2) *Main*, and 3) the *Dashboard*. In the *Admin* space, users with administrator rights can set up new projects and pipelines. Users can interact with existing projects and pipelines through the *Main* section, where the results for individual RAW files are displayed. The *Dashboard* allows users to interact with all data within a pipeline, conduct QC, detect anomalous samples, and explore abundance profiles for one or several proteins.

After a pipeline has been created, authorized users can upload one or more RAW files from the pipeline page in the *Main* interface (Figure S4). RAW files can be submitted as they are collected to get real-time feedback. Uploaded files are tracked within a PostgreSQL relational database that stores project metadata, pipeline configurations, processing status, and results. Processing tasks are executed asynchronously using Celery (https://github.com/celery/celery), with Redis (https://github.com/redis/redis-py) as the message broker. In this fashion, multiple files can be analyzed in parallel while maintaining centralized job scheduling and monitoring. Each uploaded RAW file generates a result record that includes peptide/protein identifications via MaxQuant and identification-independent scan metrics generated by RawTools. Together, these outputs provide over a hundred scan-level instrument metrics and identification-dependent statistics derived from peptide and protein identifications (Table S1). Another important aspect to consider when processing multiple files within a pipeline is False Discovery Rate (FDR) control. MaxQuant internally guarantees FDR control for each individual run. However, as the pipeline accumulates more runs, naively compiling lists of identified proteins across many experiments inflates FDR values. Here, we use the picked-protein-group FDR method to control the global FDR. This method was originally developed to work natively with MaxQuant results files and is able to combine multiple search results into a single protein group analysis.^31^

The interactive *Dashboard* is implemented with Dash and django-plotly-dash and allows users to explore results from samples processed within a given pipeline. The *Dashboard* contains three tabs: *Quality Control, Anomaly Detection*, and *Protein Explorer*. The *Quality Control* tab displays one or two user-selected QC metrics as interactive plots and allows the user to export the underlying QC table as a CSV file (Figure S5). These visualizations help users identify batch effects, technical drift, and low-quality samples. Nevertheless, identifying underlying data quality issues using a single metric can be challenging because of the high number of QC metrics, many of which are interdependent. This means that issues may be attributed to an incorrect group of metrics or to inappropriate steps in the analysis or interpretation pipeline. To address this shortcoming, we used an unsupervised multivariate Isolation Forest model^32^ that integrates multiple QC metrics and an expected outlier fraction (set to 5% in the default settings, but adjustable by the user) to identify anomalous sample runs in large cohorts. Feature contributions are interpreted using SHAP values,^33^ which help identify the QC variables most strongly associated with abnormal samples (Figure S6). Any analytical runs flagged using this approach are then highlighted in the *Quality Control* tab. It is worth noting that LAMPrEY only proposes potential outliers; the decision to exclude outliers ultimately rests with the user. The entirety of the data, including the anomaly predictions and SHAP analysis, can be accessed and downloaded from the *Quality Control* tab. Finally, the *Protein Explorer* tab allows users to examine protein abundance measurements across runs for one or more selected proteins and to inspect protein-specific trends within the cohort (Figure S7).

### Assessing LAMPrEY for longitudinal QC monitoring and biological insight

To evaluate the practical utility of LAMPrEY, we analyzed a cohort of 910 *E. faecium* isolates processed over an eight-month period. Quality control was performed for TMT11 runs acquired on an Orbitrap Fusion Lumos and processed with MaxQuant and RawTools. Across this dataset, 2,556 unique proteins were detected, with an average of 1,257 proteins quantified per isolate. The number of identified peptide sequences remained relatively stable throughout the experiment (mean value: 7,029 peptides); however, several samples exhibited markedly reduced peptide identification rates and higher fractions of missing TMT reporter values (Figure 2).

**Figure 2.**
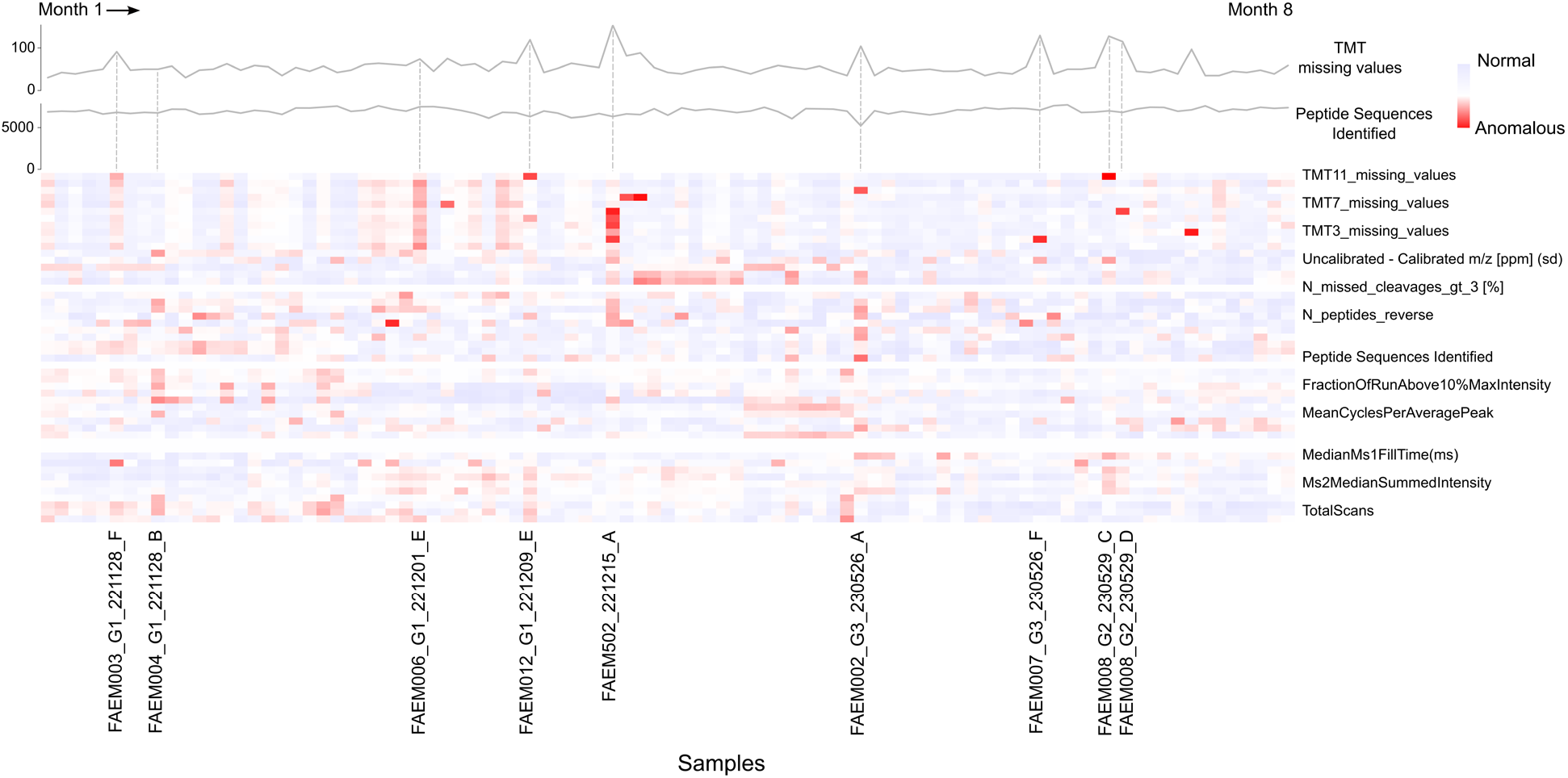
Longitudinal QC monitoring of a large proteomics cohort using LAMPrEY. QC metrics from 910 *E. faecium* isolates acquired over an eight-month period. The line plots in the upper panel show longitudinal trends for two representative metrics: the fraction of missing TMT reporter values and the number of identified peptide sequences. The heatmap summarizes multiple QC variables across runs, including TMT reporter completeness, peptide identification statistics, mass calibration accuracy, digestion quality, and instrument performance metrics. Each column corresponds to a single sample (11 TMT) and each row to a QC metric. Values are derived from a SHAP analysis, with red indicating a higher contribution to anomalous prediction. Flagged samples are listed at the bottom.

The integrated anomaly-detection algorithm in LAMPrEY flagged these samples, and SHAP analysis highlighted the TMT missing values as a higher contributor to the anomaly prediction. Based on the identification of these anomalies, we inspected the experimental setup and observed that in some instances, the liquid handler did not aspirate sufficient TMT reagent when the volume of labeling solution remaining in the tube was low. In these cases, the pipette tip likely failed to reach the bottom of the tube, resulting in incomplete labeling for a subset of samples. We were thus able to adjust the experimental approach (adding more TMT reagent) to negate this issue in all subsequent analytical runs. This example illustrates how LAMPrEY can quickly identify subtle sample-level technical anomalies within large cohorts.

LAMPrEY also facilitates the exploration of protein abundance patterns across large sample collections. For example, we examined proteins encoded by the vancomycin resistance (*van*) operon in *E. faecium* isolates (Figure 3a). The proteins VanR, VanH, VanA, VanX, and VanY are involved in remodeling peptidoglycan precursors and confer vancomycin resistance.^34^ Raw abundance profiles across isolates showed distinct abundance distributions among operon members (Figure 3a).

**Figure 3.**
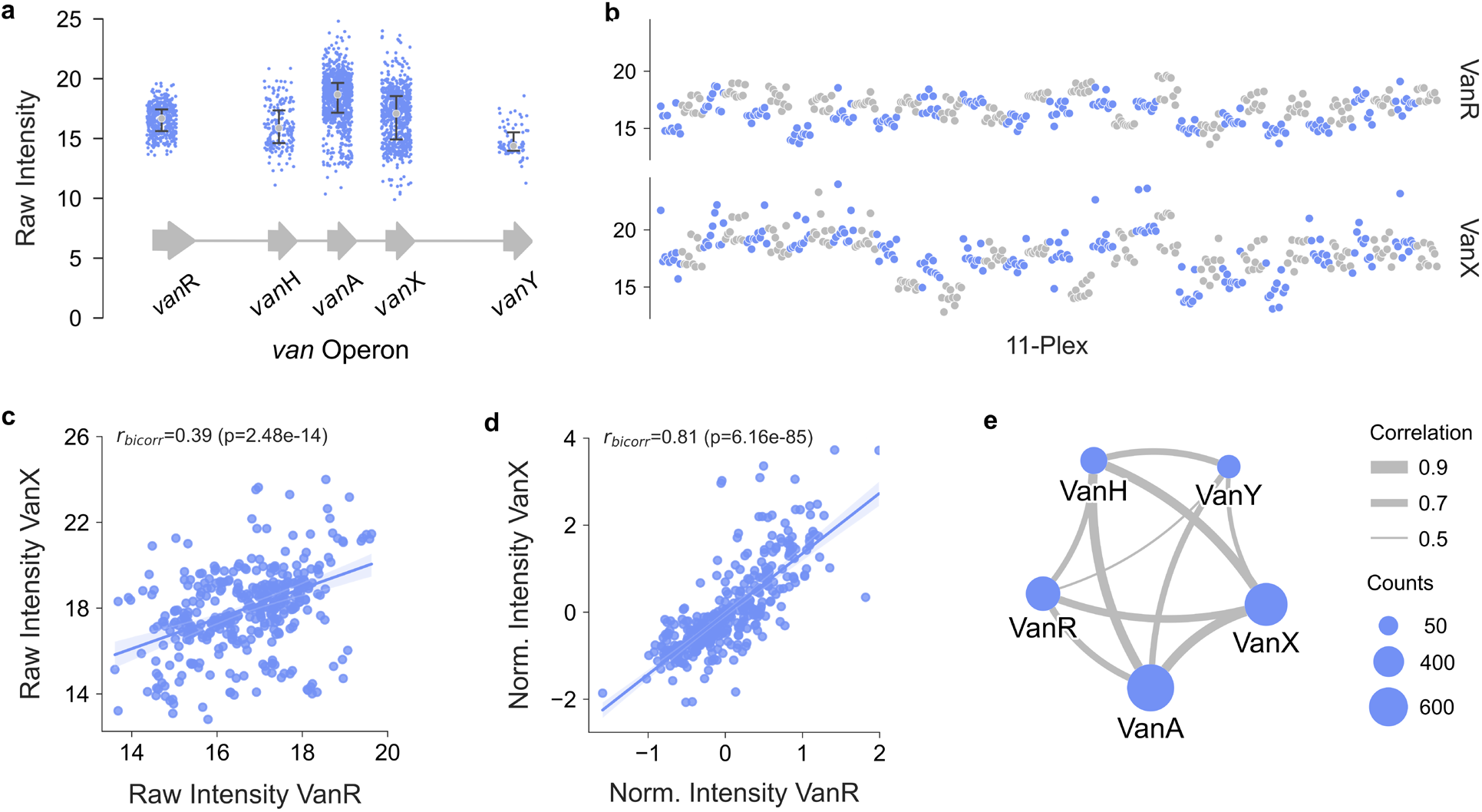
Leveraging LAMPrEY to refine analysis of protein expression from the *van* operon. a) Distribution of log_2_-transformed raw intensities for VanR, VanH, VanA, VanX, and VanY proteins across all samples. The gray dots indicate the median value of the distributions. Bars represent the interquartile range (top cap, 75^th^ percentile; bottom cap, 25^th^ percentile). The schematic at the bottom depicts the *van* operon and the order of the *van* family genes. b) log_2_-transformed raw intensities of VanR (top) and VanX (bottom) across all 11-plex batches. Alternate batches are distinguished by color. c) and d) Scatterplots of VanR versus VanX abundance with robust regression (Huber loss) fitted to raw (c) and normalized (d) intensities. Biweight midcorrelation coefficients (*r*_*bicorr*_) and corresponding p-values are indicated. e) Correlation network of VanR, VanH, VanA, VanX, and VanY. Edge thickness reflects correlation strength, and node size represents the number of quantification values for each protein.

We used LAMPrEY to examine raw intensities across this dataset and identify batch effects in individual protein intensities. For example, we observed that both VanR and VanX profiles were subject to such effects and clear differences between intensity values were present across these data (Figure 3b). These batch-to-batch differences resulted in a weak correlation between these two proteins (*r*_*bicorr*_=0.39; p-value=2.48e^-14^; Figure 3c). Normalization to a common bridge (reference) sample removed the batch-specific shifts in the data and revealed a strong linear trend between these two proteins (*r*_*bicorr*_=0.81; p-value=6.16e^-85^; Figure 3d).

We extended this analysis to all pairwise correlations and found that normalization strengthened the correlations in all cases (Tables S2 and S3); in cases with fewer observations this effect was more pronounced (*e*.*g*., VanH-VanY). Using normalized intensity values, we then built a pairwise correlation network of all the proteins encoded by the *van* operon (Figure 3e). Pairwise correlations among all members in this network were strong, with the lowest value corresponding to the VanR-VanY pair (*r*_*bicorr*_=0.62; p-value=1.63e^-06^; Figure S8) and the highest corresponding to the VanA-VanX pair (*r*_*bicorr*_=0.87; p-value=5.99e^-192^; Figure S8). These correlated abundance profiles are consistent with transcriptional co-regulation of the *van* operon. Collectively, these results illustrate the utility of LAMPrEY for identifying and mitigating batch effects in large-scale proteomic datasets.

## ABBREVIATIONS

API: application programming interface
TMT: tandem mass tag
QC: quality control
LC: liquid chromatography
MS: mass spectrometry
SHAP: shapley additive explanations.

## CONCLUSIONS AND FUTURE DIRECTIONS

Here, we introduce LAMPrEY, a Python-based platform for standardized QC monitoring and data interpretation in large-scale proteomics studies. LAMPrEY combines browser-based sample submission, automated processing, and interactive data visualization. A central feature of LAMPrEY is the application of the same processing parameters across all runs within a project. This ensures full reproducibility and enables direct comparison of QC metrics across runs. These capabilities are important for longitudinal and high-throughput experiments, where manual inspection becomes impractical. Using a cohort of 910 *E. Faecium* isolates, we demonstrate how LAMPrEY enables continuous monitoring of experimental data quality and the rapid interrogation of biologically meaningful protein abundance patterns.

The current implementation of LAMPrEY integrates RawTools and MaxQuant for data processing and supports Thermo Fisher RAW files and TMT-based quantitative workflows. In future versions, we will expand LAMPrEY to support alternative mass spectrometry formats and other MaxQuant analysis modes, such as label-free quantification, data-independent acquisition, and analysis of fractionation data. The tool we present here is open-source and available on the Lewis Research Group web page at www.lewisresearchgroup.org/software

## AUTHOR CONTRIBUTIONS

The manuscript was written through the contributions of all authors. MEV-T and SW are the project’s main authors, developers, and maintainers. MSV-T and AP assisted in developing LAMPrEY, tested its utility, and provided technical feedback. AUL and MH conducted the proteomics experiments, NIB and ELH provided constructive feedback. IAL conceived of the study, assisted with analyses, and wrote the final version of the manuscript. All authors have given approval to the final version.

## TOC

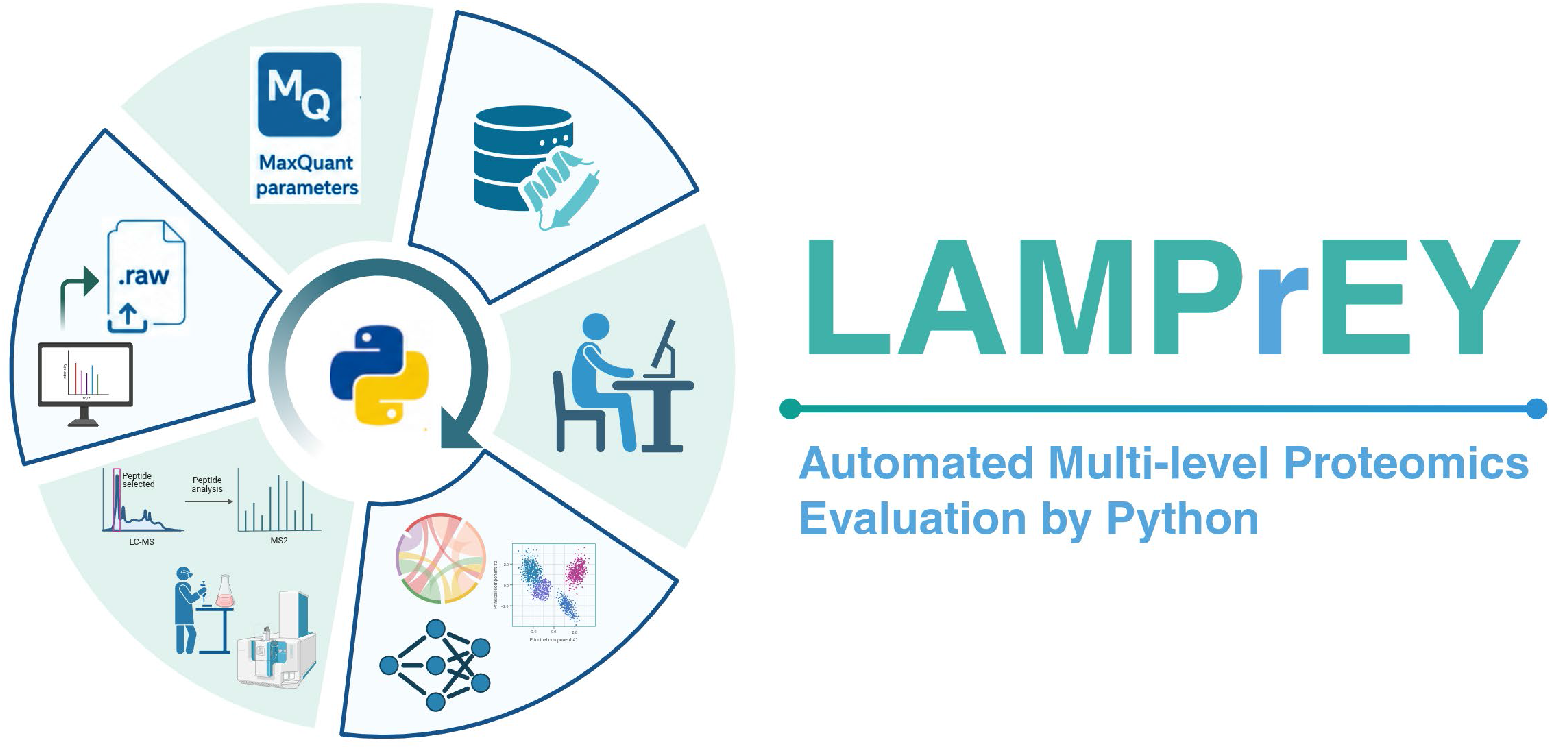

