## Supplementary Information for "LAMPrEY: a Python-based automated quality control tool for large-scale proteomics datasets"

Ian A. Lewis^1^

^1^Alberta Centre for Advanced Diagnostics (ACAD), Department of Biological Sciences, University of Calgary; Calgary, T2N 1N4, Canada

^2^ Department of Cell Biology, Harvard Medical School, Boston, Massachusetts, USA

^§^These authors contributed equally to this work

### Table S1

Table S1. Collection of ID-based and non-ID-based metrics collected in LAMPrEY.

| **Metric** | **Classification** | **Source** | **Notes** |
| --- | --- | --- | --- |
| TMT<n>_peptide_count | ID-based | App-derived | App-derived per-channel peptide counts from evidence.txt reporter intensities. |
| TMT<n>_protein_group_count | ID-based | App-derived | App-derived per-channel protein-group counts from proteinGroups.txt reporter intensities. |
| Av. Absolute Mass Deviation [mDa] | ID-based | MaxQuant QC | Average absolute mass deviation across IDs. |
| Mass Standard Deviation [mDa] | ID-based | MaxQuant QC | Mass deviation spread across IDs. |
| Missed Cleavages [%] | ID-based | MaxQuant QC | Derived as 100 - N_missed_cleavages_eq_0 [%]. |
| MS/MS Identified | ID-based | MaxQuant QC | Identified MS/MS spectra count. |
| MS/MS Identified [%] | ID-based | MaxQuant QC | Percent of submitted MS/MS spectra identified. |
| N_missed_cleavages_eq_0 [%] | ID-based | MaxQuant QC | Percent of peptides with 0 missed cleavages. |
| N_missed_cleavages_eq_1 [%] | ID-based | MaxQuant QC | Percent of peptides with 1 missed cleavage. |
| N_missed_cleavages_eq_2 [%] | ID-based | MaxQuant QC | Percent of peptides with 2 missed cleavages. |
| N_missed_cleavages_gt_3 [%] | ID-based | MaxQuant QC | Percent of peptides with more than 3 missed cleavages. |
| N_missed_cleavages_total | ID-based | MaxQuant QC | Number of peptides with at least one missed cleavage. |
| N_peptides | ID-based | MaxQuant QC | Peptide entry count. |
| N_peptides_last_amino_acid_K [%] | ID-based | MaxQuant QC | Percent of peptides ending in K. |
| N_peptides_last_amino_acid_other [%] | ID-based | MaxQuant QC | Percent of peptides ending in another residue. |
| N_peptides_last_amino_acid_R [%] | ID-based | MaxQuant QC | Percent of peptides ending in R. |
| N_peptides_potential_contaminants | ID-based | MaxQuant QC | Peptides marked as contaminants. |
| N_peptides_reverse | ID-based | MaxQuant QC | Peptides marked as reverse hits. |
| N_protein_groups | ID-based | MaxQuant QC | Protein-group count. |
| N_protein_potential_contaminants | ID-based | MaxQuant QC | Protein groups marked as contaminants. |
| N_protein_reverse_seq | ID-based | MaxQuant QC | Protein groups marked as reverse hits. |
| N_protein_true_hits | ID-based | MaxQuant QC | Protein groups excluding contaminants/reverse hits. |
| Oxidations [%] | ID-based | MaxQuant QC | Percent of peptides with oxidation annotation. |
| Peak Width (std) | ID-based | MaxQuant QC | SD of retention length over evidence rows. |
| Peak Width(ave) | ID-based | MaxQuant QC | Average retention length over evidence rows. |
| Peptide Sequences Identified | ID-based | MaxQuant QC | Unique identified peptide sequences. |
| Peptide_length_median | ID-based | MaxQuant QC | Median peptide length. |
| Peptide_msms_count_median | ID-based | MaxQuant QC | Median MS/MS count per peptide. |
| Peptide_PEP_lt_0_01 [%] | ID-based | MaxQuant QC | Percent of peptides with PEP < 0.01. |
| Peptide_PEP_median | ID-based | MaxQuant QC | Median posterior error probability. |
| Peptide_score_mean | ID-based | MaxQuant QC | Mean peptide score. |
| Peptide_score_median | ID-based | MaxQuant QC | Median peptide score. |
| Peptide_unique_groups [%] | ID-based | MaxQuant QC | Percent of peptides unique at group level. |
| Peptide_unique_proteins [%] | ID-based | MaxQuant QC | Percent of peptides unique at protein level. |
| Protein_mean_seq_cov [%] | ID-based | MaxQuant QC | Mean protein sequence coverage. |
| Protein_msms_count_median | ID-based | MaxQuant QC | Median MS/MS count per protein group. |
| Protein_peptides_median | ID-based | MaxQuant QC | Median peptide count per protein group. |
| Protein_qvalue_lt_0_01 [%] | ID-based | MaxQuant QC | Percent of protein groups with q-value < 0.01. |
| Protein_qvalue_median | ID-based | MaxQuant QC | Median protein-group q-value. |
| Protein_razor_unique_peptides_median | ID-based | MaxQuant QC | Median razor+unique peptide count per protein group. |
| Protein_score_mean | ID-based | MaxQuant QC | Mean protein-group score. |
| Protein_score_median | ID-based | MaxQuant QC | Median protein-group score. |
| Protein_unique_peptides_eq_1 [%] | ID-based | MaxQuant QC | Percent of protein groups supported by exactly one unique peptide. |
| Protein_unique_peptides_median | ID-based | MaxQuant QC | Median unique peptide count per protein group. |
| Protein_unique_seq_cov_median [%] | ID-based | MaxQuant QC | Median unique sequence coverage. |
| TMT<n>_missing_values | ID-based | MaxQuant QC | Dynamic per-channel missing-value counts from reporter intensities in proteinGroups.txt. |
| Uncalibrated - Calibrated m/z [Da] (ave) | ID-based | MaxQuant QC | Average absolute mass calibration shift over evidence rows. |
| Uncalibrated - Calibrated m/z [Da] (sd) | ID-based | MaxQuant QC | SD of absolute mass calibration shift over evidence rows. |
| Uncalibrated - Calibrated m/z [ppm] (ave) | ID-based | MaxQuant QC | Average mass calibration shift over evidence rows. |
| Uncalibrated - Calibrated m/z [ppm] (sd) | ID-based | MaxQuant QC | SD of mass calibration shift over evidence rows. |
| CutoffDecoyScore(0.05FDR) | ID-based | RawTools QC | Decoy score cutoff at 5% FDR. |
| DigestionEfficiency | ID-based | RawTools QC | Digestion efficiency computed from identified peptides/PSMs. |
| IdentificationRate(IDs/Ms2Scan) | ID-based | RawTools QC | Identification rate normalized by MS2 scans. |
| MedianPeptideScore | ID-based | RawTools QC | Median peptide identification score. |
| MissedCleavageRate(/PSM) | ID-based | RawTools QC | Missed-cleavage rate across PSMs. |
| NumberOfPSMs | ID-based | RawTools QC | Number of peptide-spectrum matches. |
| NumberOfUniquePeptides | ID-based | RawTools QC | Number of unique identified peptides. |
| PsmChargeRatio3to2 | ID-based | RawTools QC | Charge-state ratio among identified PSMs. |
| PsmChargeRatio4to2 | ID-based | RawTools QC | Charge-state ratio among identified PSMs. |
| Mean_parent_int_frac | Non-ID-based | MaxQuant QC | Mean parent intensity fraction from msmsScans.txt. |
| MS | Non-ID-based | MaxQuant QC | MaxQuant summary scan count. |
| MS/MS | Non-ID-based | MaxQuant QC | MaxQuant summary MS/MS scan count. |
| MS/MS Submitted | Non-ID-based | MaxQuant QC | Spectra submitted for searching. |
| MS3 | Non-ID-based | MaxQuant QC | MaxQuant summary MS3 scan count. |
| DateAcquired | Non-ID-based | RawTools QC | RawTools QC export timestamp for the run. |
| ExperimentMsOrder | Non-ID-based | RawTools QC | Acquisition method metadata. |
| FractionOfRunAbove10%MaxIntensity | Non-ID-based | RawTools QC | Fraction of run above 10% max intensity. |
| Instrument | Non-ID-based | RawTools QC | Instrument metadata. |
| MeanCyclesPerAveragePeak | Non-ID-based | RawTools QC | Average cycles sampled per peak. |
| MeanDutyCycle(s) | Non-ID-based | RawTools QC | Average duty cycle. |
| MeanMs2TriggerRate(/Ms1Scan) | Non-ID-based | RawTools QC | Average MS2 triggers per MS1 scan. |
| MedianAsymmetryAt10%H | Non-ID-based | RawTools QC | Median peak asymmetry at 10% height. |
| MedianAsymmetryAt50%H | Non-ID-based | RawTools QC | Median peak asymmetry at 50% height. |
| MedianMassDrift(ppm) | Non-ID-based | RawTools QC | Median mass drift. |
| MedianMs1FillTime(ms) | Non-ID-based | RawTools QC | Median MS1 fill time. |
| MedianMs1IsolationInterference | Non-ID-based | RawTools QC | Median MS1 isolation interference. |
| MedianMs2FillTime(ms) | Non-ID-based | RawTools QC | Median MS2 fill time. |
| MedianMs2PeakFractionConsumingTop80PercentTotalIntensity | Non-ID-based | RawTools QC | MS2 peak-complexity proxy from RawTools QC. |
| MedianMs3FillTime(ms) | Non-ID-based | RawTools QC | Median MS3 fill time. |
| MedianPeakWidthAt10%H(s) | Non-ID-based | RawTools QC | Median chromatographic peak width at 10% height. |
| MedianPeakWidthAt50%H(s) | Non-ID-based | RawTools QC | Median chromatographic peak width at 50% height. |
| MedianPrecursorIntensity | Non-ID-based | RawTools QC | Median precursor intensity. |
| Ms1Analyzer | Non-ID-based | RawTools QC | MS1 analyzer metadata. |
| Ms1MedianSummedIntensity | Non-ID-based | RawTools QC | Median summed MS1 intensity. |
| Ms1ScanRate(/s) | Non-ID-based | RawTools QC | MS1 acquisition rate. |
| Ms2Analyzer | Non-ID-based | RawTools QC | MS2 analyzer metadata. |
| Ms2MedianSummedIntensity | Non-ID-based | RawTools QC | Median summed MS2 intensity. |
| Ms2ScanRate(/s) | Non-ID-based | RawTools QC | MS2 acquisition rate. |
| Ms3Analyzer | Non-ID-based | RawTools QC | MS3 analyzer metadata. |
| Ms3ScanRate(/s) | Non-ID-based | RawTools QC | MS3 acquisition rate. |
| NumEsiInstabilityFlags | Non-ID-based | RawTools QC | Count of electrospray instability flags. |
| NumMs1Scans | Non-ID-based | RawTools QC | MS1 scan count. |
| NumMs2Scans | Non-ID-based | RawTools QC | MS2 scan count. |
| NumMs3Scans | Non-ID-based | RawTools QC | MS3 scan count. |
| PeakCapacity | Non-ID-based | RawTools QC | Peak capacity estimate. |
| RawFile | Non-ID-based | RawTools QC | Raw data file name/path emitted by RawTools. |
| SearchParameters | Non-ID-based | RawTools QC | Search-method metadata recorded by RawTools. |
| TimeAfterLastExceedanceOf10%MaxIntensity | Non-ID-based | RawTools QC | Time after chromatogram falls below 10% max intensity. |
| TimeBeforeFirstExceedanceOf10%MaxIntensity | Non-ID-based | RawTools QC | Time before chromatogram rises above 10% max intensity. |
| TotalAnalysisTime(min) | Non-ID-based | RawTools QC | Total run duration. |
| TotalScans | Non-ID-based | RawTools QC | Total scans acquired. |

### Table S2

Table S2. Pairwise biweight midcorrelation calculated using raw intensity profiles for proteins encoded in the *van* operon.

| P1 | P2 | n | *r_bicorr_* | CI95 | p_unc | p_corr | power |
| --- | --- | --- | --- | --- | --- | --- | --- |
| VanA | VanH | 170 | 0.72 | [0.64, 0.78] | 2.18E-28 | 5.46E-28 | 1 |
| VanA | VanX | 615 | 0.79 | [0.75, 0.81] | 3.98E-130 | 3.98E-129 | 1 |
| VanA | VanY | 70 | 0.71 | [0.57, 0.81] | 6.42E-12 | 9.17E-12 | 1 |
| VanA | VanR | 380 | 0.56 | [0.49, 0.63] | 5.02E-33 | 1.67E-32 | 1 |
| VanH | VanX | 170 | 0.8 | [0.74, 0.85] | 2.22E-39 | 1.11E-38 | 1 |
| VanH | VanY | 20 | -0.23 | [-0.61, 0.24] | 0.33113 | 0.33113 | 0.165223 |
| VanH | VanR | 150 | 0.56 | [0.44, 0.66] | 1.05E-13 | 1.76E-13 | 1 |
| VanX | VanY | 70 | 0.45 | [0.24, 0.62] | 8.49E-05 | 0.000106 | 0.980073 |
| VanX | VanR | 360 | 0.39 | [0.30, 0.47] | 2.48E-14 | 4.96E-14 | 1 |
| VanY | VanR | 50 | 0.22 | [-0.06, 0.47] | 0.121671 | 0.13519 | 0.34438 |

**P1**: protein 1

**P2**: protein 2

**n**: number of observations considered for the correlation

***r_bicorr_***: biweight midcorrelation coefficient

**CI95**: 95% parametric confidence intervals

**p_unc**: uncorrected p-values

**p_corr**: corrected p-values with Benjamini/Hochberg FDR correction

**power**: achieved power of the test (= 1 - type II error)

### Table S3

Table S3. Pairwise biweight midcorrelation calculated using normalized intensity profiles for proteins encoded in the *van* operon.

| P1 | P2 | n | *r_bicorr_* | CI95 | p_unc | p_corr | power |
| --- | --- | --- | --- | --- | --- | --- | --- |
| VanA | VanH | 170 | 0.87 | [0.83, 0.90] | 1.39E-53 | 3.47E-53 | 1 |
| VanA | VanX | 615 | 0.87 | [0.85, 0.89] | 5.99E-193 | 5.99E-192 | 1 |
| VanA | VanY | 70 | 0.76 | [0.64, 0.85] | 1.59E-14 | 2.27E-14 | 1 |
| VanA | VanR | 380 | 0.78 | [0.74, 0.82] | 4.98E-80 | 1.66E-79 | 1 |
| VanH | VanX | 170 | 0.86 | [0.82, 0.90] | 1.99E-51 | 3.99E-51 | 1 |
| VanH | VanY | 20 | 0.78 | [0.51, 0.91] | 5.77E-05 | 5.77E-05 | 0.991503 |
| VanH | VanR | 150 | 0.75 | [0.67, 0.81] | 5.80E-28 | 9.67E-28 | 1 |
| VanX | VanY | 70 | 0.7 | [0.56, 0.80] | 1.31E-11 | 1.64E-11 | 1 |
| VanX | VanR | 360 | 0.81 | [0.77, 0.84] | 6.16E-85 | 3.08E-84 | 1 |
| VanY | VanR | 50 | 0.62 | [0.41, 0.77] | 1.47E-06 | 1.63E-06 | 0.998918 |

**P1**: protein 1

**P2**: protein 2

**n**: number of observations considered for the correlation

***r_bicorr_***: biweight midcorrelation coefficient

**CI95**: 95% parametric confidence intervals

**p_unc**: uncorrected p-values

**p_corr**: corrected p-values with Benjamini/Hochberg FDR correction

**power**: achieved power of the test (= 1 - type II error)

### Figure S1

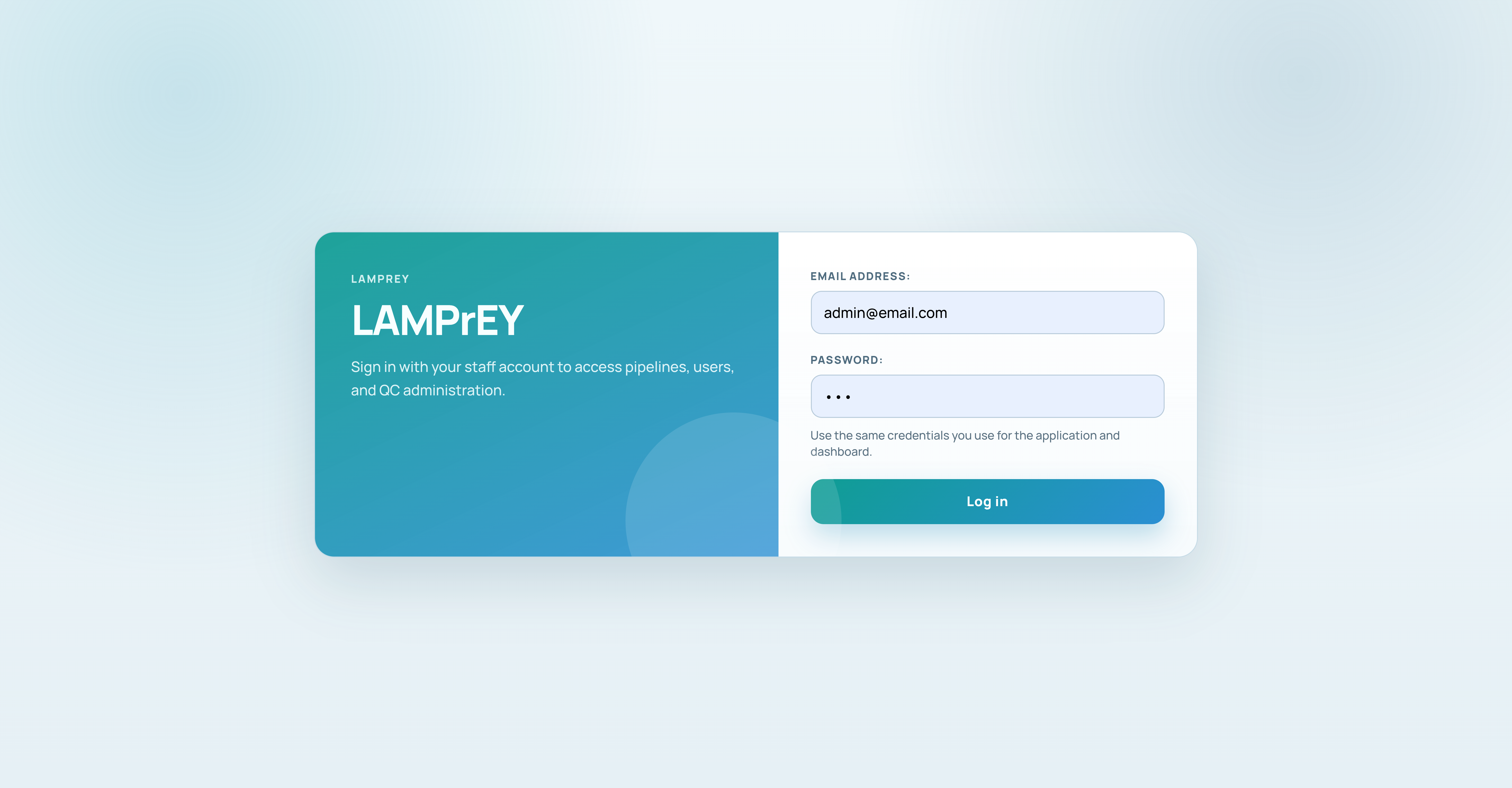

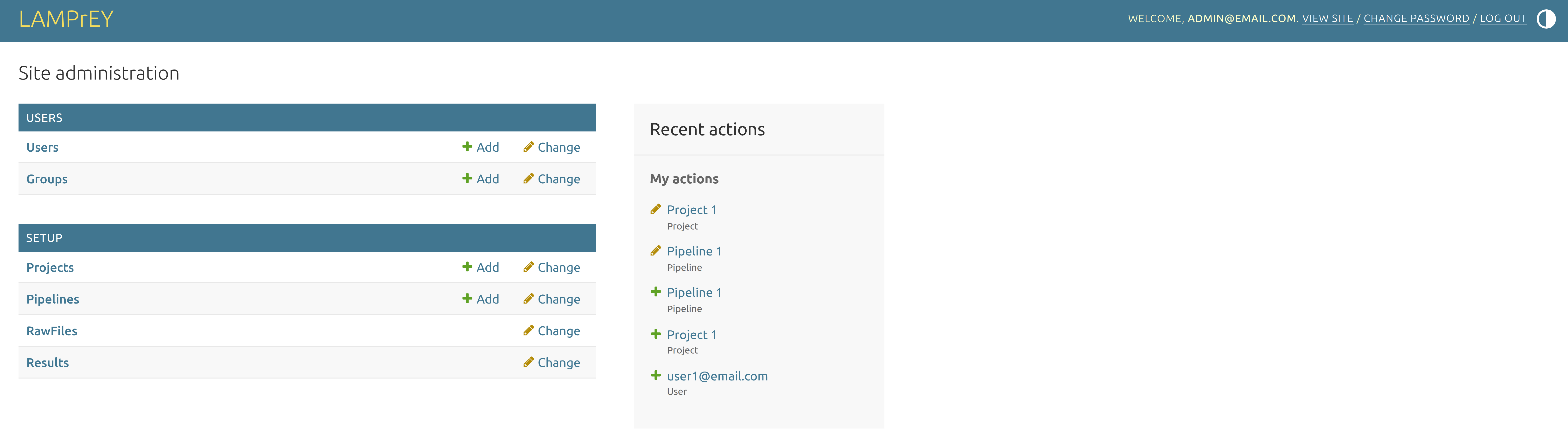

Figure S1. Admin panel in LAMPrEY. The admin section is password-protected and manages projects, pipelines, raw files, and all results.

### Figure S2

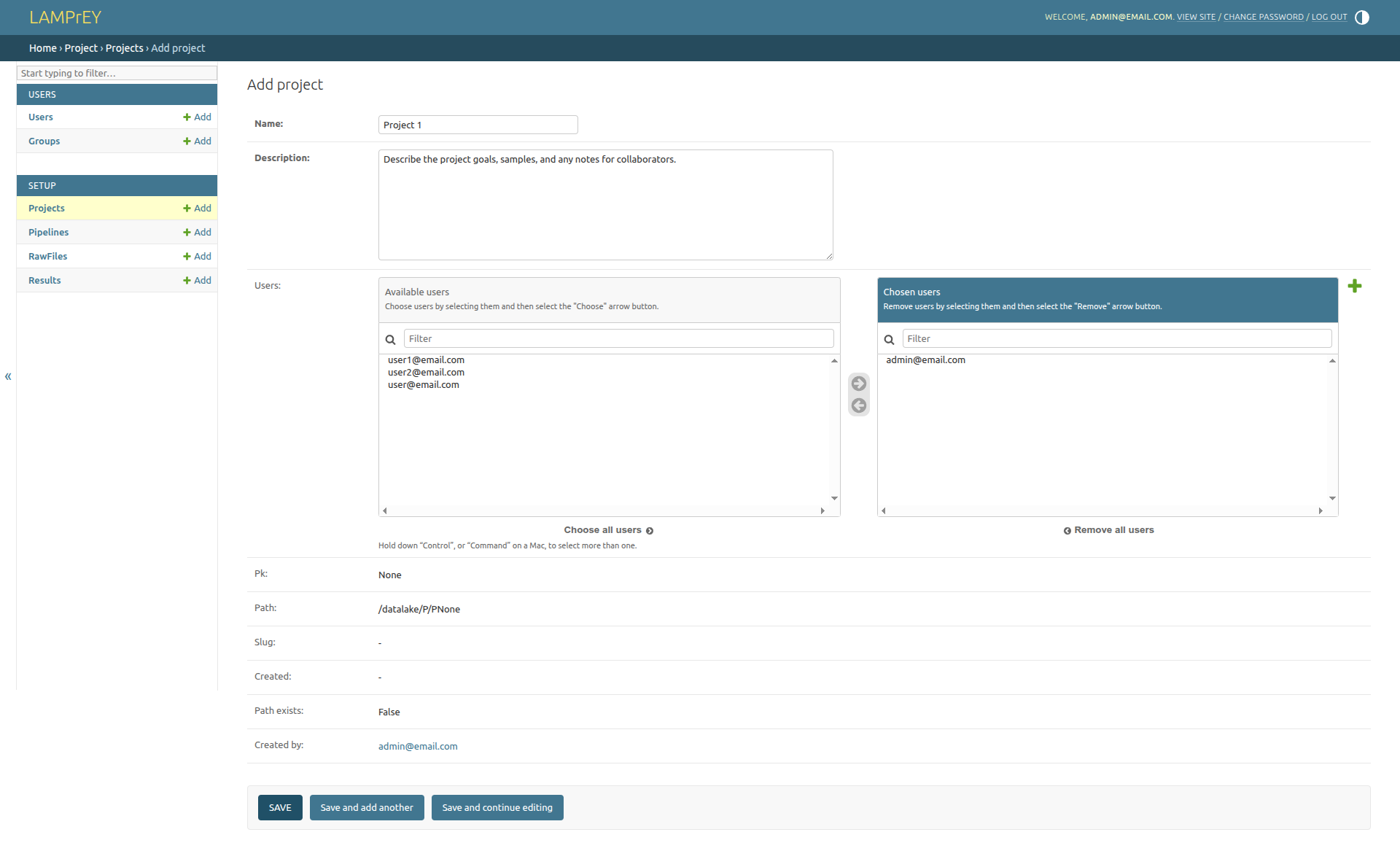

Figure S2. Section “Add project” within the admin panel. Projects are independent workspaces and can contain one or multiple pipelines.

### Figure S3

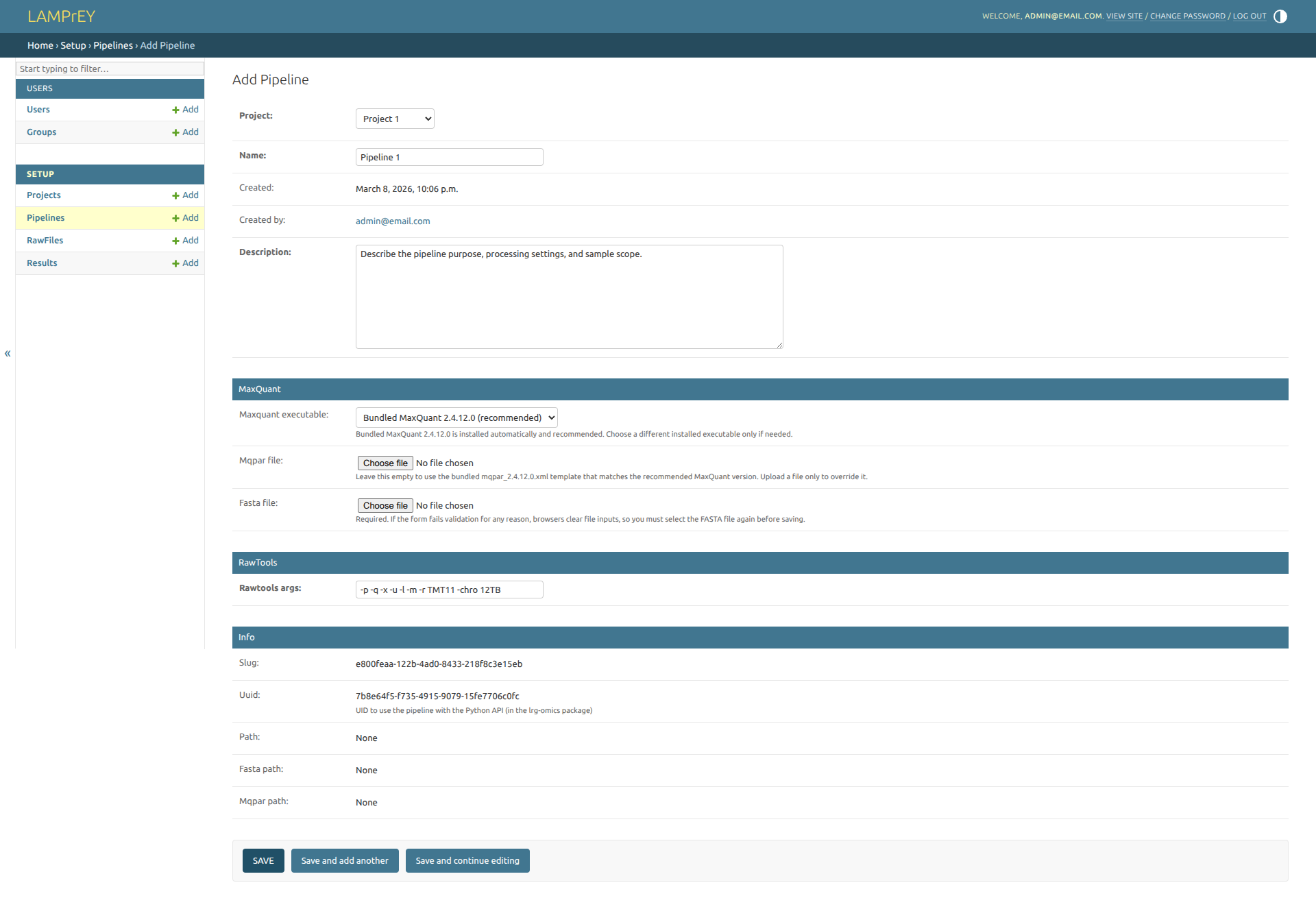

Figure S3. Section “Add pipeline” within the admin panel. Pipelines are assigned to a certain project and require a working MaxQuant executable, a suitable mqpar.xml file, and a FASTA file with the protein database.

### Figure S4

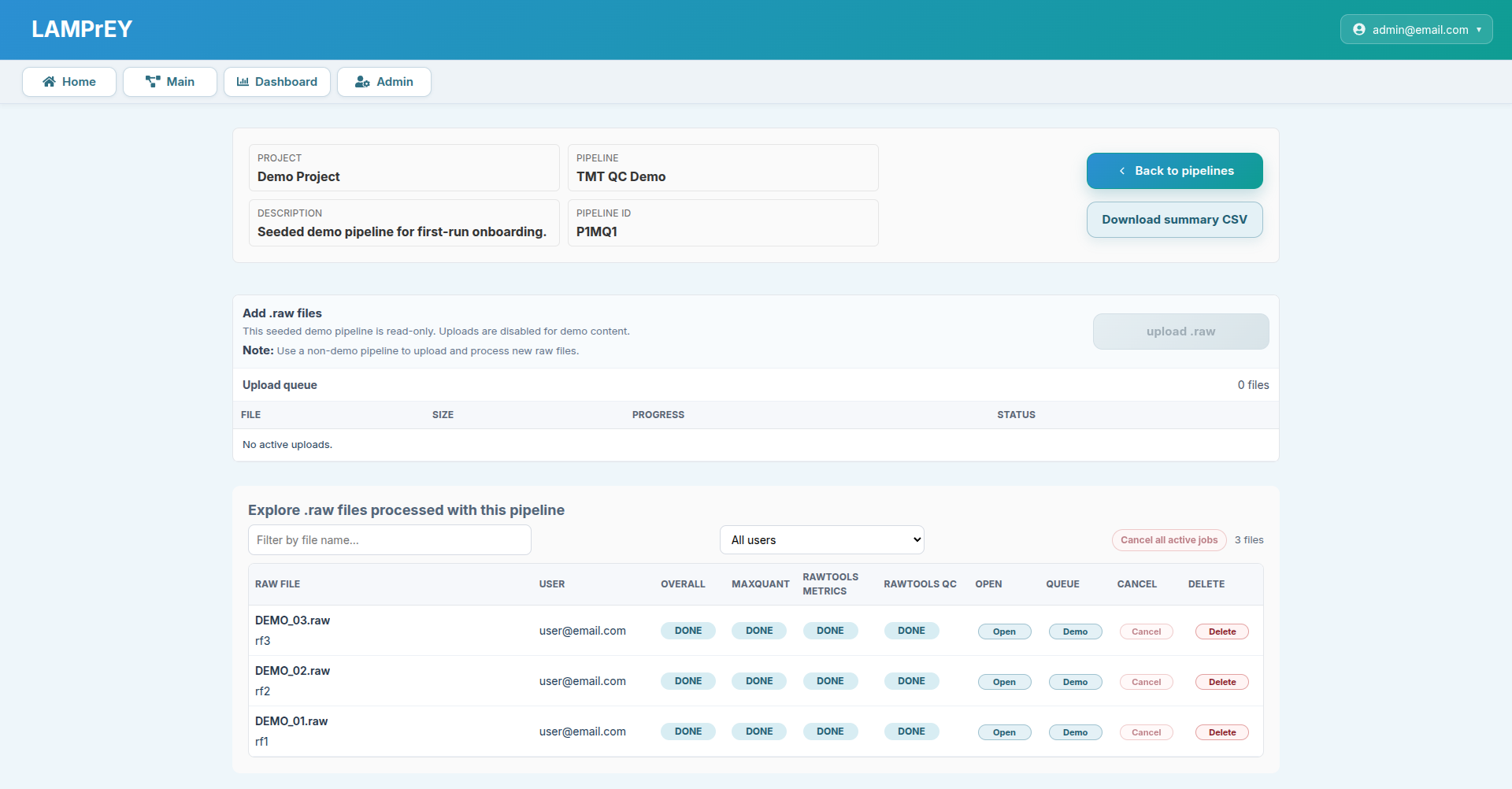

Figure S4. Main section in LAMPrEY. This section allows users to verify the number of submitted RAW files, monitor queue status, and review the results of processed files. The RAW files listed in the image here are preloaded demo datasets that are included for demonstration purposes.

### Figure S5

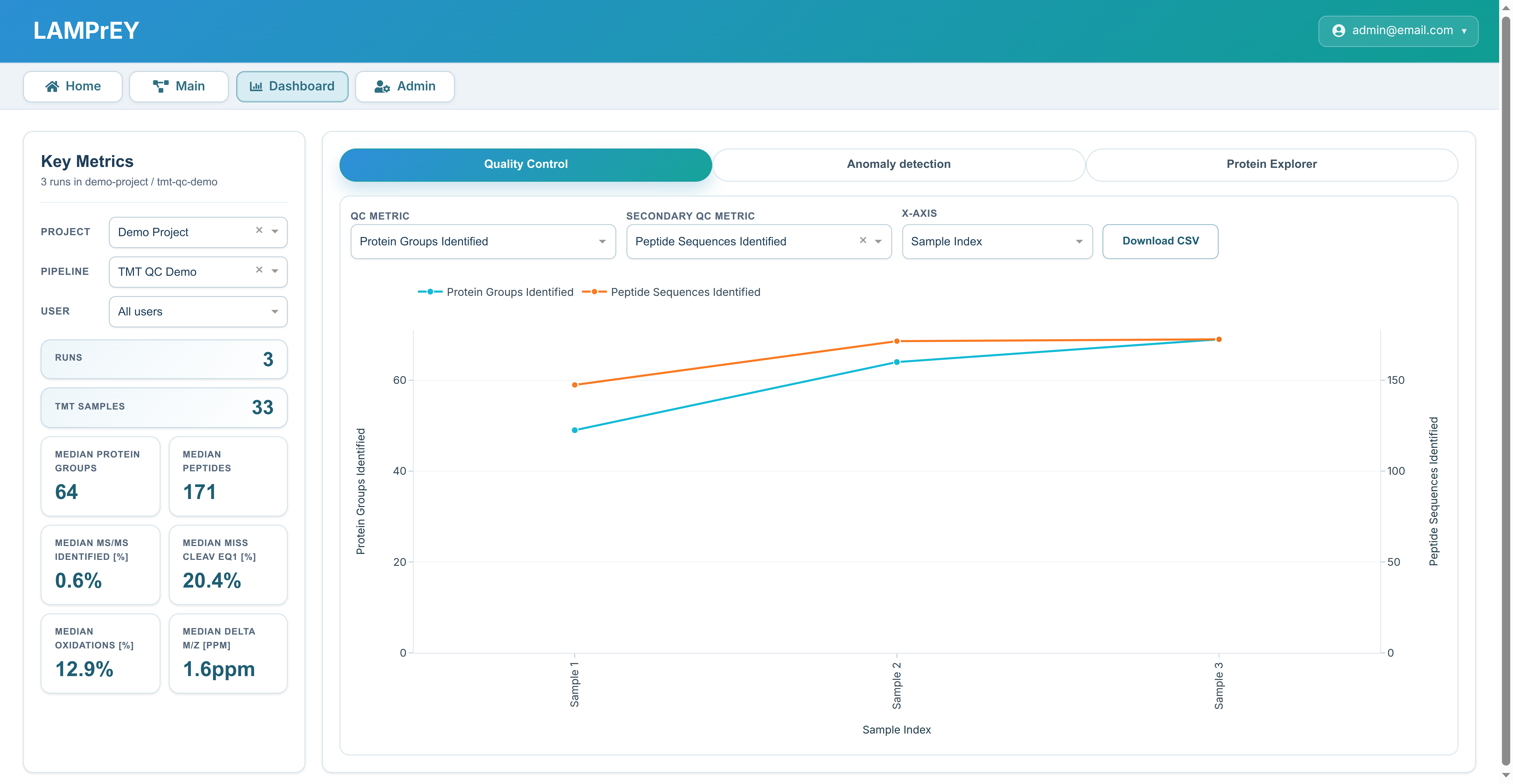

Figure S5. Quality Control tab on the Dashboard. Selected QC metrics are displayed as interactive plots and can be exported as a CSV file.

### Figure S6

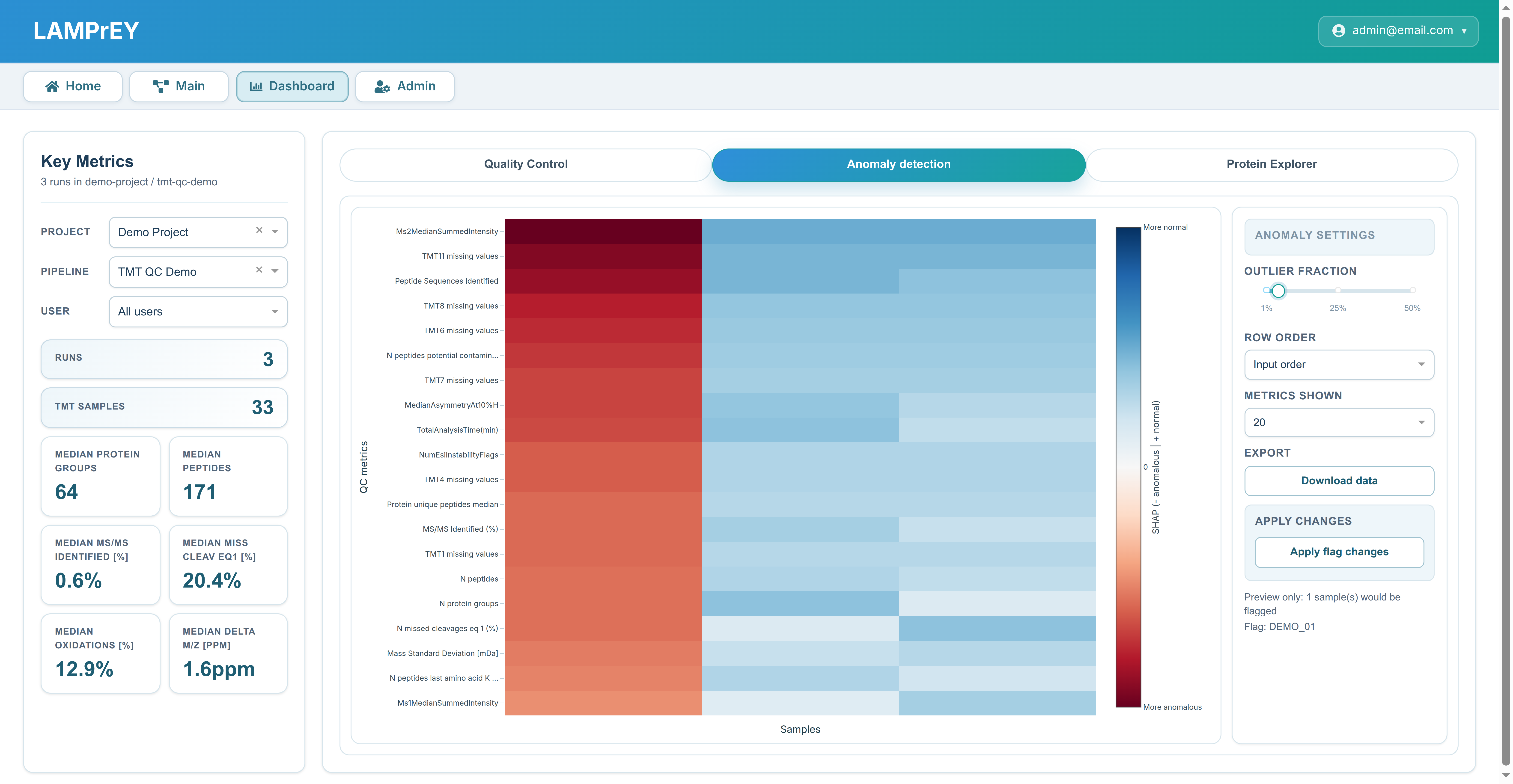
**Figure S**6**. Anomaly Detection tab on the Dashboard.** The Isolation Forest method is used to flag anomalous samples. Heatmap values are derived from SHAP analysis and are used to identify QC metrics most strongly associated with anomalous samples.

### Figure S7

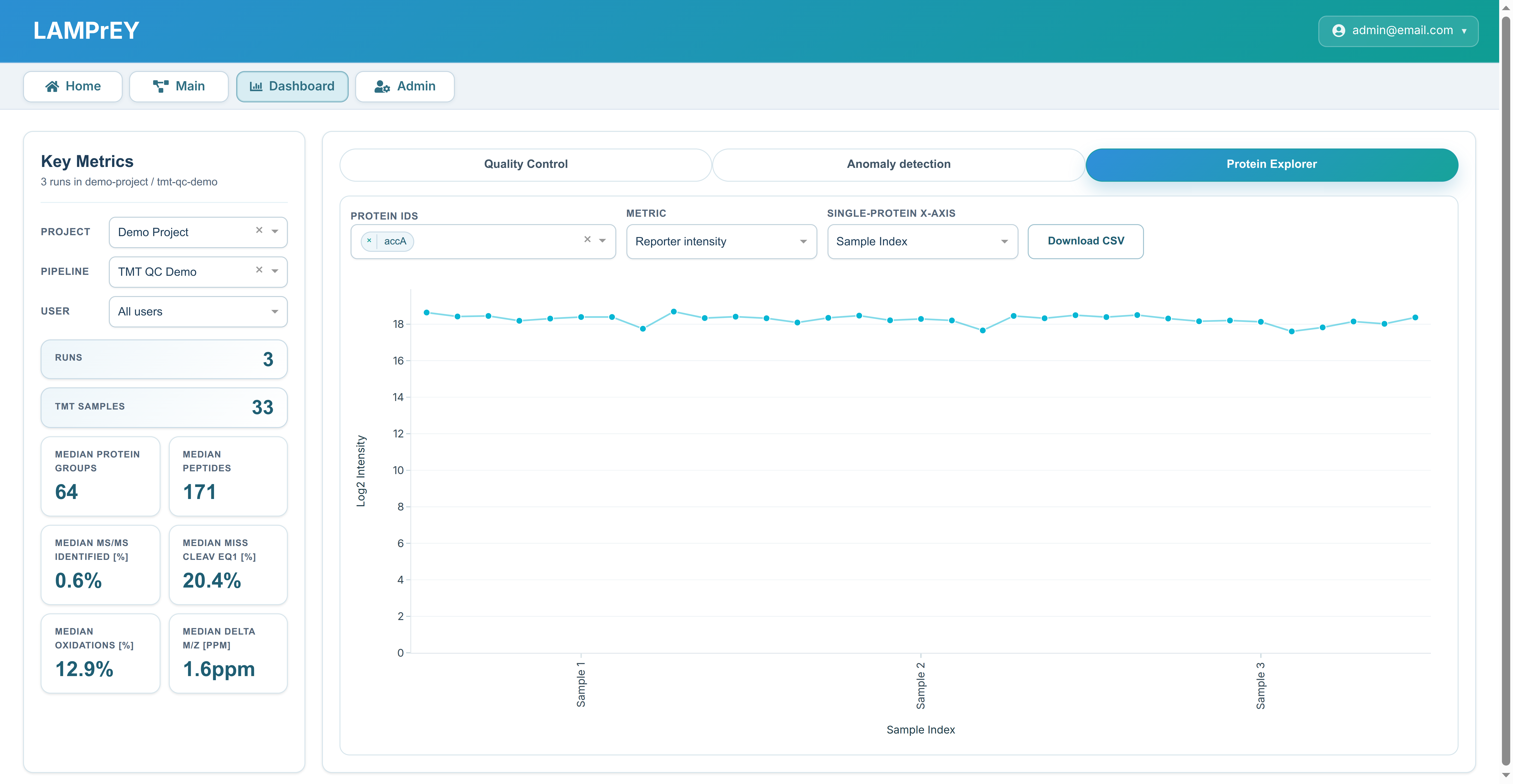

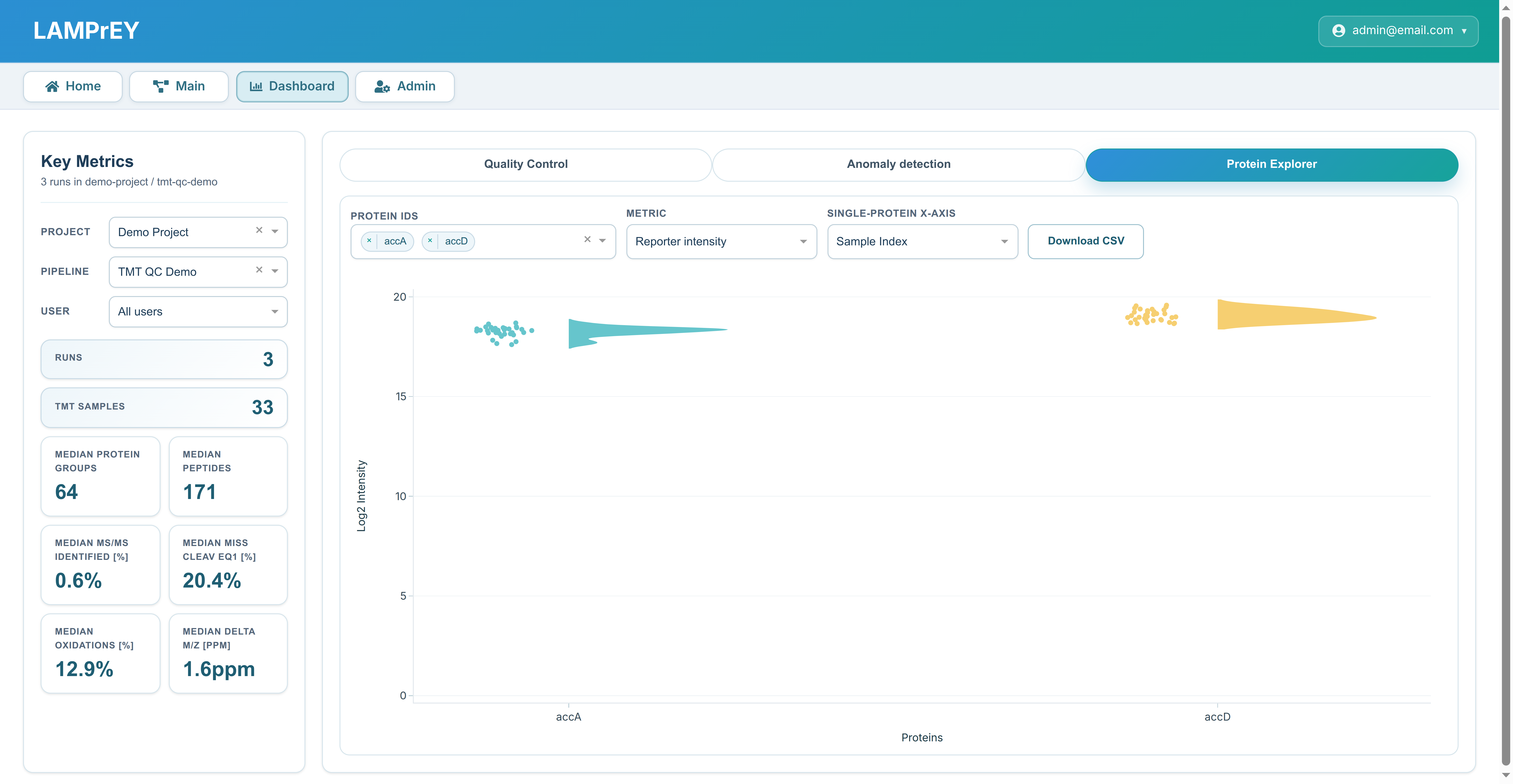

Figure S7. Protein Explorer tab on the Dashboard. Log2 intensities can be retrieved for a single protein (top panel) or multiple proteins (bottom panel). The top panel shows intensities for Acetyl-CoA carboxylase carboxyl transferase subunit alpha. The bottom panel shows intensities for Acetyl-CoA carboxylase carboxyl transferase subunits alpha and beta. In both cases, intensities are shown across 33 TMT samples (three runs, 11 TMT samples each).

### Figure S8

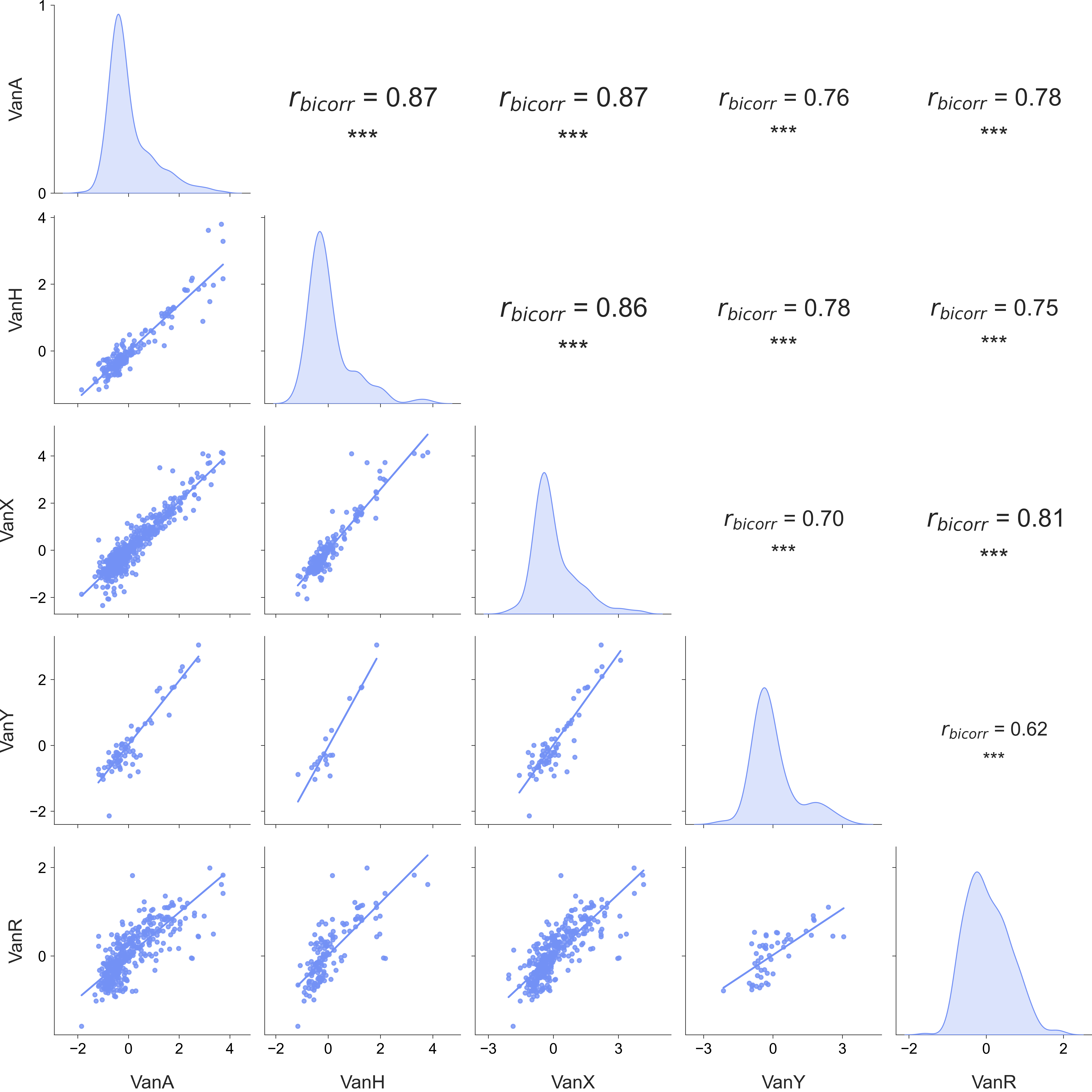

Figure S8. Biweight midcorrelation among VanA, VanH, VanX, VanY, and VanR normalized abundance profiles. Scatterplots with robust regression lines fitted using the Huber loss are shown in the lower triangle, kernel density distributions are shown on the diagonal, and biweight midcorrelation coefficients (*r_bicorr_*) with significance levels are shown in the upper triangle. Asterisks indicate significance thresholds (* p < 0.05, ** p < 0.01, *** p < 0.001; ns, not significant).
